# G-quadruplexes sequester free heme in living cells

**DOI:** 10.1101/652297

**Authors:** Lucas T. Gray, Emilia Puig Lombardi, Daniela Verga, Corinne Guetta, Alain Nicolas, Marie-Paule Teulade-Fichou, Arturo Londoño-Vallejo, Nancy Maizels

**Affiliations:** Department of Biochemistry, University of Washington, 1959 NE Pacific St., Seattle, WA 98195, USA; Allen Institute for Brain Science, Seattle, Washington, 98109 USA; Institut Curie, Centre de Recherche, CNRS-UMR3244, PSL Research University, 75005 Paris, France; Institut Curie, Centre de Recherche, CNRS-UMR9187, INSERM-U1196, PSL Research University, Sorbonne Universités, 91405 Orsay, France; Department of Immunology, University of Washington, 1959 NE Pacific St., Seattle, WA 98195, USA

**Keywords:** G-quadruplex, heme, iron homeostasis, gene expression, oxidative damage, genomic stability

## Abstract

Heme is an essential cofactor for many enzymes, but free heme is toxic and its levels are tightly regulated. G-quadruplexes bind heme avidly *in vitro*, raising the possibility that they may sequester heme *in vivo*. If so, then treatment that displaces heme from quadruplexes is predicted to induce expression of genes involved in iron and heme homeostasis. Here we show that PhenDC3, a G-quadruplex ligand structurally unrelated to heme, displaces quadruplex-bound heme *in vitro* and alters transcription in cultured human cells, up-regulating genes that support heme degradation and iron homeostasis, and most strikingly causing a 30-fold induction of heme oxidase 1, the key enzyme in heme degradation. We propose that G-quadruplexes sequester heme to protect cells from the pathophysiological consequences of free heme. This identifies a new function for G-quadruplexes and a new mechanism for protection of cells from heme.

## Introduction

G-quadruplexes are nucleic acid structures composed of stacked ‘G-quartets’ [Gellert M et al. 1962; Sen D and Gilbert W 1988], planar arrays of four hydrogen-bonded guanines (**Figure 1A**) connected by loops composed of other nucleotides. Quadruplexes form at sequences bearing the G4 motif, consisting of four or more runs of guanines punctuated by other nucleotides: G_≥3_N_x_G_≥3_N_x_G_≥3_N_x_G_≥3_. In the human genome, there are nearly 400,000 G4 motifs allowing loops of ≤ 7 nt [Huppert JL and Balasubramanian S 2005], and more than 700,000 allowing loops up to 12 nt [Maizels N and Gray LT 2013]. G4 motifs are dispersed throughout the genome and considerably enriched at promoters, where they flank the transcription start site (TSS) and cluster at the 5’ end of the first intron [Eddy J and Maizels N 2008; Eddy J and Maizels N 2009]. Analyses of individual G4 motifs have identified potential regulatory roles of quadruplexes in replication, transcription, splicing, and translation [Bugaut A and Balasubramanian S 2012; Tarsounas M and Tijsterman M 2013; Maizels N 2015; Valton AL and Prioleau MN 2016; Svikovic S and Sale JE 2016; Weldon C et al. 2017]. G4 motifs can be sites of replication arrest and they are closely associated with genetic and epigenetic stability [Lemmens B et al. 2015; Maizels N 2015; Svikovic S and Sale JE 2016], but G4 motifs that are thermodynamically stable and therefore genetically unstable have been purged from most genomes [Puig Lombardi E et al. 2019]. This makes the abundance of G4 motifs especially puzzling, and suggests that they may play useful roles in cell physiology that have yet to be identified.

**Figure 1.**
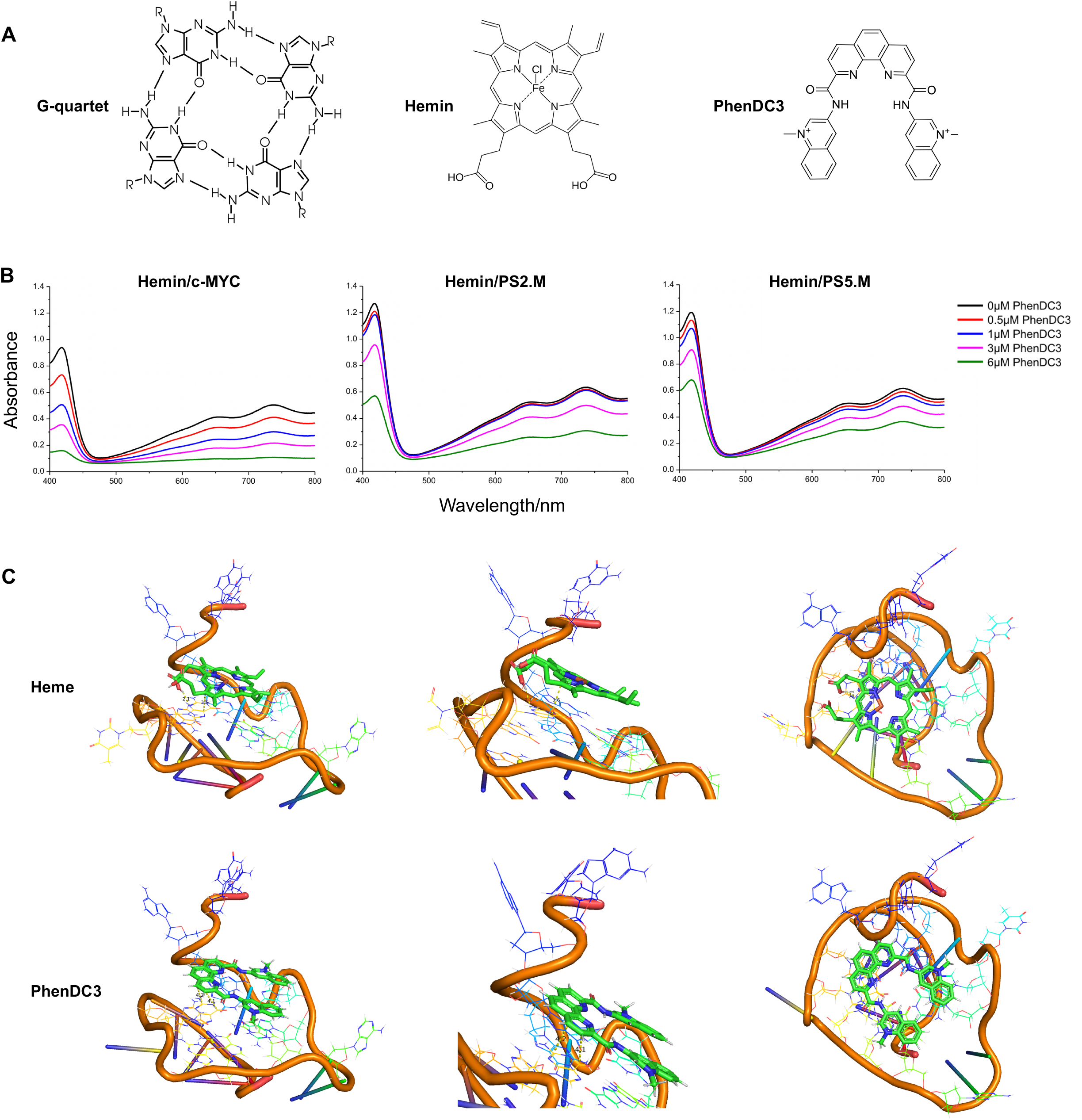
PhenDC3 displaces hemin from three different G-quadruplex structures. **(A)** Structures of a G-quartet, Fe(III)-heme and PhenDC3 (structures are not size-scaled). **(B)** Absorption spectra analyzing H_2_O_2_-dependent oxidation of ABTS^2-^ to ABTS^1-^ by G4 DNAzyme in the presence of the G4 ligand PhenDC3 (0-6 μM). G4 DNAzymes assayed were complexes of hemin with quadruplexes formed from the indicated oligonucleotides: hemin/c-MYC, hemin/PS2.M and hemin/PS5.M. **(C)** Above: structure of Fe(III) hemin (PDB accession: HEM) docked on the c-MYC promoter DNA G-quadruplex, shown as (from left to right) a side view, a zoomed side view and a top view. The distance from the loop to the iron at the center of the heme is ~3.6 Å (middle panel, dotted yellow line). Below: model view of the solution structure of the G4 ligand PhenDC3 bound to the c-MYC G-quadruplex (PDB coordinates accession: 2MGN), shown as (from left to right) a side view, a zoomed side view and a top view. For the PyMOL visualizations, we used default atom colors. In particular, hemin iron is dark orange (RGB code triples R=0.87843137; G=0.400000000 and B=0.200000000); refer to https://pymolwiki.org/index.php/Color_Values).

G-quadruplexes bind avidly to heme *in vitro* (Kd ≈ 10 nM range), as first shown more than 20 years ago [Li Y and Sen D 1996] and subsequently pursued in many contexts (reviewed in [Canale TD and Sen D 2017]). Indeed, the size, planarity and hydrophobicity of the protoporphyrin IX scaffold are well suited for stacking with the terminal G-quartets while, in addition, guanine or cytosine residues may provide axial coordination to the ferric ion [Li T et al. 2009], altogether making heme a privileged ligand for G-quadruplexes. This binding mode has been confirmed by molecular dynamics [Nasab MG et al. 2017] and docking studies [Poon LC et al. 2011]. Heme is an essential cofactor in a great variety of processes involving oxidation, oxygen transport and electron transport, and just as protein enzymes use heme as a prosthetic group, G-rich nucleic acid aptamers can use the electron transfer capabilities of a bound heme to support peroxidase reactions [Poon LC et al. 2011]. The natural affinity of quadruplexes for heme suggested that quadruplexes may bind to heme *in vivo*, enabling use of heme as a cofactor for nucleic acid-catalyzed reactions [Poon LC et al. 2011]. However, free heme is potentially toxic even at low concentrations, as its redox-active iron can catalyze formation of harmful reactive oxygen species and promoting oxidative stress [Chiabrando D et al. 2014; Ponka P et al. 2017]. This raised the possibility that sequestration of heme by quadruplexes might protect cells and the genome from oxidative damage by limiting free heme levels while ensuring ready availability of this key cofactor.

Here we experimentally test the possibility that quadruplexes sequester heme in living cells. While it would be very difficult to directly quantify very small amounts of intracellular heme bound to quadruplexes, free heme initiates its own elimination by inducing well-documented changes in gene expression, including induction of transcription of the genes encoding heme oxidase 1 (HMOX1), the key enzyme in heme degradation, and ferritin (FTH1 and FTL1), the iron storage protein [Kappas A and Drummond GS 1986; Hamamura RS et al. 2007; Ghosh S et al. 2011]. Moreover, a number of ligands that interact with G-quadruplexes have been shown to readily displace quadruplex-bound heme *in vitro* [Kong DM et al. 2008] and are predicted to similarly displace heme bound to quadruplexes *in vivo*. We therefore designed an experimental strategy that would enable us to detect release of heme from G-quadruplexes by using RNA-Seq analysis to query the effect of treatment with a G-quadruplex ligand predicted to displace sequestered heme on expression of genes involved in heme catabolism and iron homeostasis. We chose to use the well-characterized G4 ligand PhenDC3, a bisquinolinium phenanthroline derivative structurally unrelated to heme (**Figure 1A**) [De Cian A et al. 2007; Piazza A et al. 2010; Halder R et al. 2012; Chung WJ et al. 2014; Perriaud L et al. 2014; Lista MJ et al 2017]. PhenDC3 is an ideal compound for genome-wide studies of quadruplex function *in vivo*, because it readily enters the nucleus [Lefebvre J et al. 2017]; it discriminates well between quadruplex and duplex DNA; and it exhibits a very high affinity (nanomolar Kd) for quadruplexes, without pronounced selectivity for any specific quadruplex structure. This lack of quadruplex selectivity can be attributed to the extensive overlap of PhenDC3 with the guanines in a G-quartet, in the absence of minor groove interactions characteristic of some G4 ligands, which enables PhenDC3 to bind with high affinity to the external G-quartets of a quadruplex [Chung WJ et al. 2014].

In this work, we validate displacement of G4-bound heme by PhenDC3 *in vitro* and we show, by RNA-seq analysis, that PhenDC3 treatment significantly up-regulates pathways involved in heme metabolism and iron homeostasis in human HT1080 fibrosarcoma cells, most strikingly causing a 30-fold induction of HMOX1, the chief enzyme in the heme degradation pathway. A search of publicly available transcriptome datasets showed that a similar restricted subset of genes is induced by treatment with hemin [Ghosh S et al. 2011] and with a structurally unrelated G4 ligand, the anthraquinone derivative AQ1 [Zorzan E et al. 2018]. These findings identify a new mechanism for regulation of free heme levels in living cells. They also identify an unanticipated potential function for quadruplexes in maintaining genomic integrity that takes advantage of the abundance of G4 motifs and their dispersion throughout the genome.

## Results and Discussion

### PhenDC3 can displace quadruplex-bound hemin

G-quadruplexes can associate with hemin (Fe(III)-heme) to form G4 DNAzymes which can catalyze peroxidation reactions [Li Y and Sen D 1996; Travascio P et al. 1998], and displacement of hemin by a G4 ligand inhibits this catalytic activity. To evaluate the ability of PhenDC3 to displace hemin from a G-quadruplex, we used the chromogenic substrate 2,2’-azino-bis(3-ethylbenzothiazoline-6-sulfonic acid) (ABTS^2-^) as a probe. H2O2-mediated oxidation of ABTS^2-^ produces the corresponding radical ABTS^·-^ which exhibits a characteristic green signal readily detected by naked-eye and upon absorbance measurement. We tested PhenDC3-mediated displacement of hemin from three different G-quadruplex structures, one formed from an oligonucleotide representing a 22 nt G4 motif identified at the c-MYC protooncogene [Simonsson T et al. 1998], which folds to form a parallel structure [Ambrus A et al. 2005]; and two others formed by oligonucleotides, PS2.M (18 nt) and PS5.M (24 nt), both of which have been reported to fold to form antiparallel structures in the experimental conditions employed and to have strong catalytic activity once bound to hemin [Travascio P et al. 1998; Travascio P et al. 2001] (see *Materials and Methods*). A strong decrease of the absorption spectra intensity of ABTS^·-^ between 500 nm and 800 nm was observed at increasing concentrations of PhenDC3 (**Figure 1B**), as predicted if displacement of hemin from the quadruplex structures reduces the catalytic activity of the hemin/G-quadruplex DNAzymes. This trend was observed for quadruplexes formed from all three sequences tested. PhenDC3 seems to be able to displace hemin more efficiently from the c-MYC sequence (**Figure 1B, left**), with catalytic activity is almost completely suppressed after the addition of 2 molar equivalents (6 μM) of PhenDC3. More efficient displacement may reflect a higher affinity of PhenDC3 for the c-MYC quadruplex, combined with a lower affinity of hemin towards this structure, shown by the slightly reduced enzymatic activity observed in the absence of PhenDC3 (ABTS^·-^ maximum absorption at 740 nm is lower if compared to the spectra reported for PS2.M and PS5.M). The c-MYC quadruplex forms a parallel structure [Ambrus A et al. 2005], while the proposed structure for both PS2.M and PS5.M is antiparallel [Travascio P et al. 2001]), and this may affect hemin binding. Overall, these results show that hemin is released free in solution upon PhenDC3 binding to the G-quadruplex structure.

**Figure 1C** shows the predicted location for the lowest energy simulated structure obtained upon docking of heme to a high-resolution NMR structure (PDB accession: 2MGN) [Chung WJ et al. 2014] of the c-MYC promoter quadruplex (see *Materials and Methods*). The heme stacks upon a loop guanine, which in turn stacks upon the G-quartets (**Figure 1C, above**). PhenDC3 similarly binds with high affinity to an external G-quartet (**Figure 1C, below**). Thus, heme and PhenDC3 are predicted to have similar favored binding sites and lie within similar G-quartet to ligand distances (axial distance ~3.6 Å for heme and ~4.1 Å for PhenDC3; **Figure 1C**).

### Response to PhenDC3 correlates with G4 motif frequency

To profile the transcriptome of PhenDC3-treated cells, we used RNA-seq analysis to characterize gene expression in HT1080 cells, a human fibrosarcoma cell line with stable diploid karyotype. Control experiments showed that treatment with 20 μM PhenDC3 for 48 hr had only a modest effect on cell viability (**Figure 2A**) or cell cycle (**Figure 2B; Figure S1**). Whole-cell RNA was isolated from cells which were untreated or treated with PhenDC3 under these conditions, as three biological replicates (**Figure S1**). mRNA was sequenced and reads were mapped to GENCODE genes in the hg19 human reference genome (see *Materials and Methods*). G4 motifs were then located using a regular expression matching algorithm that allowed loops of 1 to 12 nt in length, G_≥3_N_1-12_G_≥3_N_1-12_G_≥3_N_1-12_G_≥3_, N={A,T,C,G}. This search criterion identifies 722,264 G4 motifs in the human genome, approximately twice as many as a search for motifs with 1 to 7 nt loops [Huppert JL and Balasubramanian S 2005], and was used because the longer loop length better captures G4 motifs associated with biological processes [Gray LT et al. 2014].

**Figure 2.**
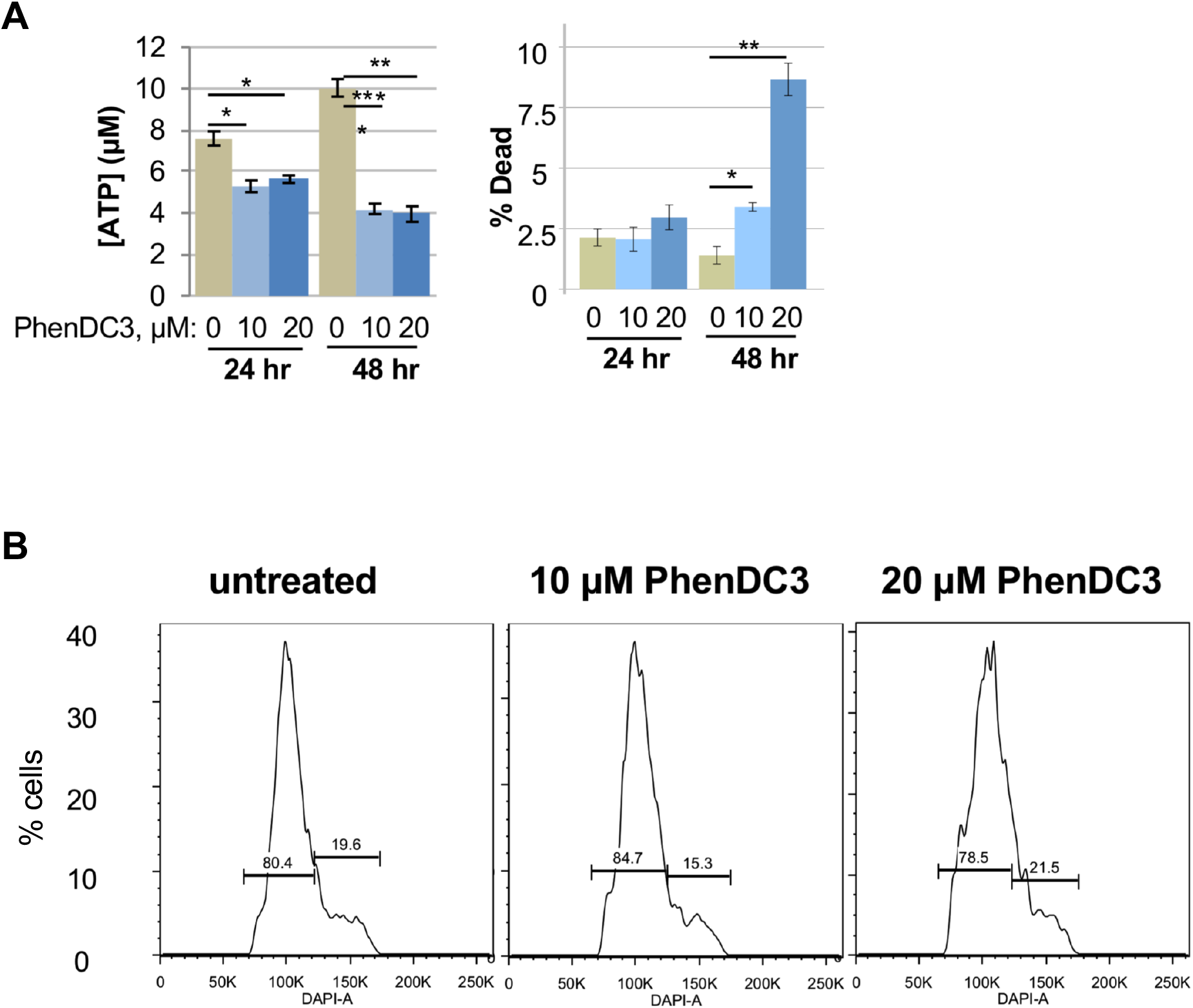
Effects of PhenDC3 on cell viability and cell cycle. **(A)** Viability of cells treated with DMSO carrier alone (1%) or 10 or 20 μM PhenDC3 in DMSO carrier as measured by ATP concentration (left) and live/dead staining (right). Error bars indicate standard deviation among triplicate samples. *, p < 0.01; **, p < 0.001; ***, p < 0.0001. **(B)** DAPI cell cycle profiles for populations treated as in panel A for 48 hr.

PhenDC3 treatment caused significant changes in the expression profiles of HT1080 cells, with samples treated with 20 μM PhenDC3 or no PhenDC3 clustering uniquely upon hierarchical classification (**Figure 3A; Figure S2**). A total of 1,745 genes exhibited ≥2-fold differential expression in response to PhenDC3 treatment, among them 1,014 up-regulated and 731 down-regulated genes (FDR<0.01) (**Figure 3B; Table S1**). Mapping G4 motif frequencies of these ≥2-fold differentially expressed genes (2xDEGs) established strong correlations between transcript abundance and enrichment or depletion of G4 motifs near the promoter and at the 5’ end of intron 1 (FDR<0.001; **Figure 3C**). This correlation was assessed by empirical calculation of false discovery rates based on random sampling from all expressed genes (see *Materials and Methods*). Furthermore, the proportion of genes carrying at least one G4 motif in the promoter region is significantly higher among 2xDEGs than genes which were unaffected (FDR>0.01) by PhenDC3 treatment (two-proportions z-test *P* = 0.0025 for down-regulated genes, *P* = 0.0032 for up-regulated genes; **Figure 3D**). In particular, up-regulated genes carry significantly more G4 sequences than any other gene set (**Figure 3D**).

**Figure 3.**
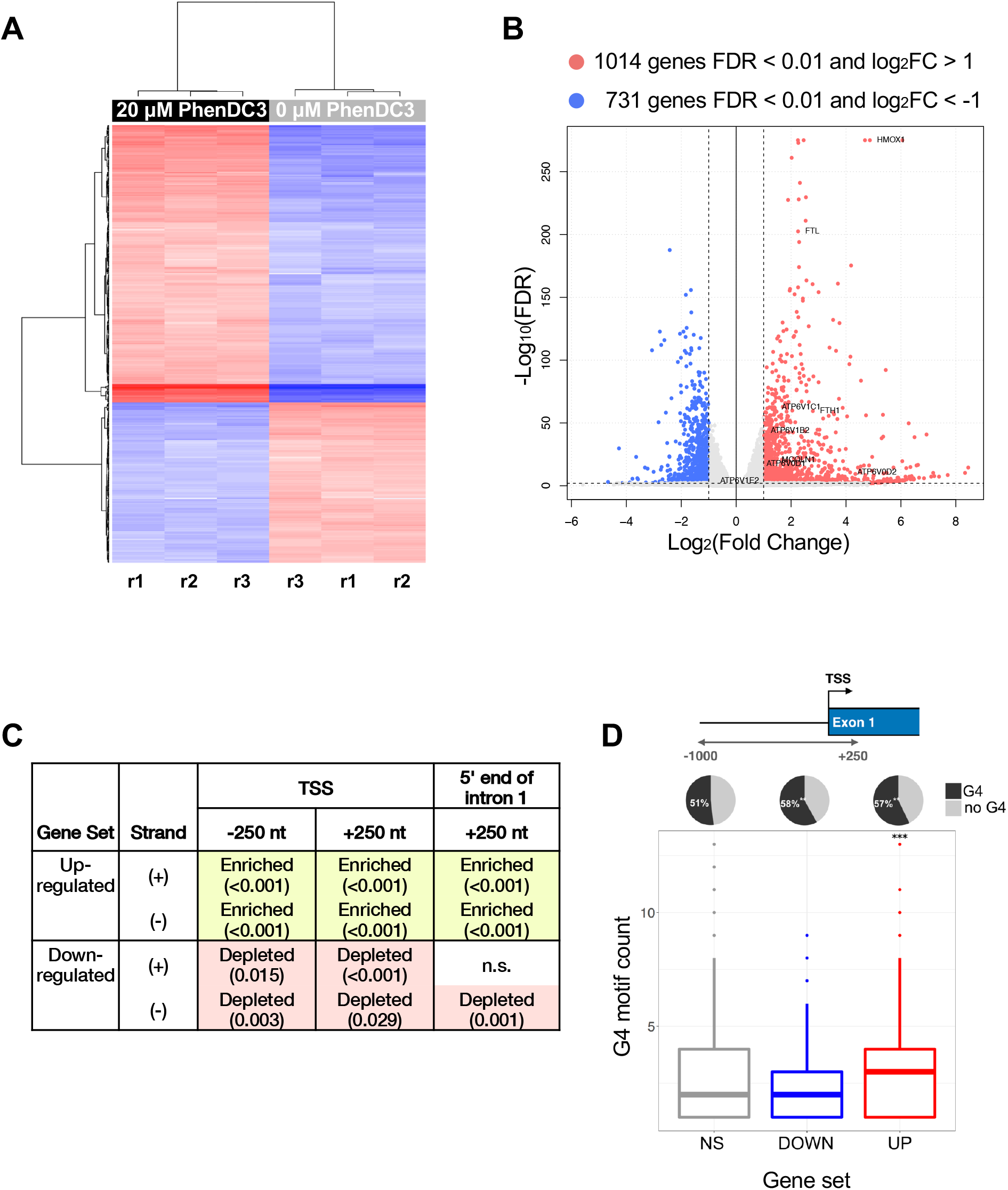
PhenDC3 treatment induces significant transcriptional changes in HT1080 fibrosarcoma cells. **(A)** Supervised clustering of log-transformed gene counts obtained from three independent RNA-seq experiments performed in HT1080 cells treated with PhenDC3 (20 μM) for 48 hours. Top scoring 500 genes are shown. **(B)** Volcano plot of genes differentially expressed (FDR < 0.01) in cells treated or untreated with PhenDC3 (20 μM). A subset of the genes significantly upregulated (Log_2_(Fold Change) > 0) is shown. **(C)** G4 ([G_3+_N_1-12_]_3+_G_3+_) enrichment and depletion statistics for 250 nt windows flanking the TSS and downstream of the 5’ end of first introns. Enrichment (green) and depletion (red) are relative to 1000 sets of randomly selected gene drawn from a pool of all genes expressed at detectable levels. FDR values in parentheses. FDR > 5% (0.05) is considered non-significant (n.s, no highlight). **(D)** G4 ([G_3+_N_1-12_]_3+_G_3+_) detection in promoter regions (1000 nt upstream and 250 nt downstream of TSS) of 2xDEGs (red, up-regulated; blue, down-regulated genes) and unaltered genes (gray, FDR>0.01). Pie charts show the number of genes carrying at least one G4 motif (black) or no G4 motif (gray) within promoter regions. Boxplots: ***, pairwise Wilcoxon tests adj*P* < 0.001 compared to NS; Pie charts: **, two-proportion z-tests *P* < 0.01.

We identified a very similar pattern of enrichment and depletion of G4 motifs upon application of our analytical and mapping techniques to a previous array-based dataset that determined the response to PhenDC3 treatment of HeLa cells [Halder R et al. 2012], a highly transformed human cervical cancer cell line (**Figure S3**). These results confirm the premise that PhenDC3 treatment accentuates the physiological functions of G4 motifs, and support the correlation between transcript abundance and enrichment of G4 motifs near the promoter (**Figure 3C**).

PhenDC3 appears to interact with both DNA and RNA quadruplexes to perturb gene expression. Positive correlations were evident with up-regulation by PhenDC3 and G4 motifs at regions immediately flanking the TSS (**Figure 3**), likely to reflect PhenDC3 interactions with DNA; and with differential splicing and enrichment of G4 motifs on the non-transcribed strand at the 5’ end of intron 1 (FDR=0.004; **Figure S4**), likely to reflect PhenDC3 interactions with pre-mRNA.

### G4 ligands induce pathways associated with heme degradation and iron homeostasis

The top ten REACTOME pathways that correlated with up-regulated 2xDEGs are shown in **Figure 4A** (full list is reported in **Table S2**). Two of the top pathways are directly related to heme and iron homeostasis (*‘Iron uptake and transport’, ‘Transferrin endocytocis and recycling’*); and three other pathways are related via the circadian rhythm/CLOCK pathway, for which the key regulator is a transcription factor (REV-ERB) that binds heme as an essential cofactor [Raghuram S et al. 2007]. Gene set enrichment analysis of up-regulated 2xDEGs also correlated with two other specific pathways, associated with immune responses and nerve growth factor (NGF) signaling. Heme binds to proteins to modulate their function, acting as a signaling molecule in a variety of biological processes, and it has been reported that these include innate immunity [Figueiredo RT et al. 2007; Dutra F and Bozza MT 2014] and neuronal survival [reviewed in Smith AG et al. 2011], which may contribute to these correlations. To observe the interaction between the up-regulated 2xDEGs, we performed network analysis which showed how these genes clustered mainly into the four afore-mentioned pathways (**Figure 4A**). The network was generated by mapping the significant genes to the STRING protein-protein interactome [Szklarczyk D et al. 2015] and applying a search algorithm to identify first-order neighbors for each of the mapped genes (see *Materials and Methods*). We generated a highly-connected first-order network (1,301 nodes and 4,038 edges), indicating concerted action of the up-regulated genes in key cellular processes involving heme.

**Figure 4.**
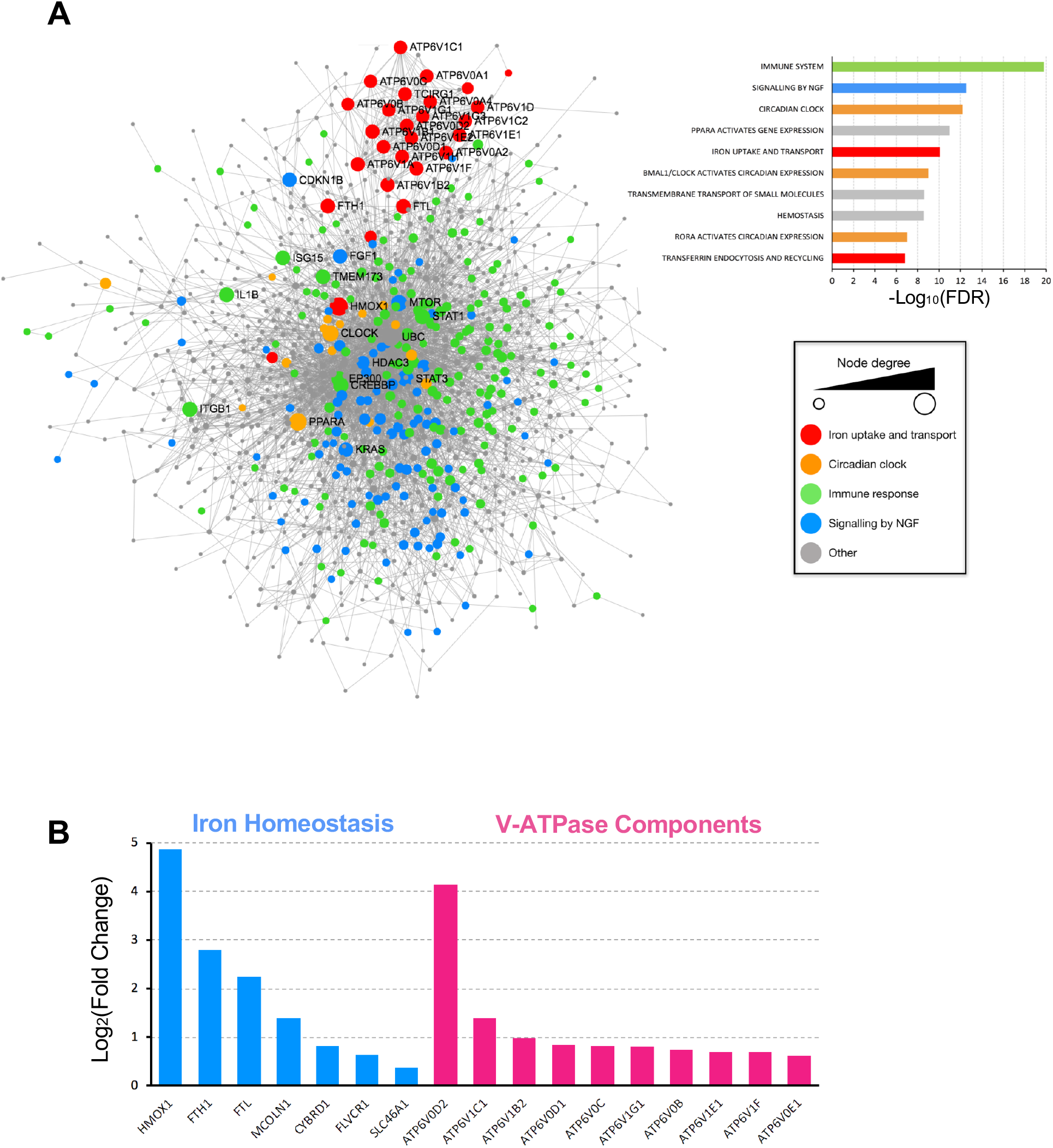
PhenDC3 treatment activates genes involved in heme and iron homeostasis. **(A)** Minimal first-order protein-protein interaction networks between up-regulated 2xDEGs. Node size is proportional to the number of its connections with other nodes (n = 1,301 nodes). The barplot shows REACTOME pathway-based enrichment analysis of genes strongly up-regulated (Log_2_(Fold Change) > 1; FDR < 0.01) in treated (+PhenDC3) relative to untreated samples. The top 10 results correlated with upregulated genes are shown. FDR, GSEA false discovery rate q-value. **(B)** Log_2_(Fold Changes) for common genes involved in iron homeostasis REACTOME pathways.

Especially striking was the 30-fold (log_2_FC=4.9) up-regulation of HMOX1 (**Figure 4B; Table S3**), which cleaves excess heme to initiate its degradation. In addition, FTH1 and FTL, which encode the two chains of the ferritin iron-storage complex, were up-regulated 5-to 7-fold. Highly significant up-regulation also was evident for the mucolipin channel protein (MCOLN1), a component of the cationic channel for Fe^++^ transport; a heme transporter (SLC46A1) and a heme exporter (FLVCR1). PhenDC3 treatment also induced transcription of several components of the vacuolar ATPase (V-ATPase) complex involved in iron and copper transport (**Figure 4B; Table S3**).

The altered regulation of iron-related pathways prompted us to analyze the expression of genes that encode iron-sulfur proteins. These proteins undergo iron-regulated biogenesis in the mitochondrion [Lill R 2009], where G4 ligands can localize due to their lipophilic cationic features and also upon binding with quadruplexes in mitochondrial DNA [Huang WC et al. 2015]. Of 26 genes surveyed, 3 were modestly but significantly up-regulated in response to PhenDC3 treatment, and 15 were down-regulated (**Figure S5; Table S4**). The only gene exhibiting more than 2-fold down-regulation was the G4 helicase, RTEL1 (log_2_FC=-1.1). These results suggest that the primary effect of PhenDC3 may be mediated by heme release, rather than catabolism of heme that releases iron.

To provide further support for the notion that induction of genes involved in heme catabolism and iron homeostasis by PhenDC3 depends upon heme release, we addressed the possibility that, despite the structural dissimilarity of these compounds (**Figure 1A**), treatment with exogenous hemin and PhenDC3 might affect transcriptional regulation of a largely overlapping set of genes. The responses to PhenDC3 and to hemin were compared by determining the overlap of the sets of genes significantly up-regulated (log_2_(fold change) > 0.5 and FDR < 0.05) in two cultured endothelial cell lines (PMVEC and PAEC) treated with hemin [Ghosh S et al. 2011] and in HT1080 cells treated with PhenDC3. The overlap included four genes, HMOX1, FTH1, GCLM and NQO1 (**Figure 5A**). These genes all share a common regulator, NRF2, a transcriptional activator that is stabilized upon binding to heme to stimulate transcription of genes involved in heme and iron metabolism, including HMOX1, FTH1 and FTL; and in antioxidant pathways, including GCLM, the rate-limiting enzyme of glutathione synthesis; and NQO1, a member of the NAD(P)H dehydrogenase family [reviewed in Tonelli C et al. 2018]. The overlap between genes induced by treatment with exogenous hemin or PhenDC3 supports the hypothesis that PhenDC3 displaces heme, and raises the possibility that heme-dependent activation of NRF2 may contribute to the transcriptional response to PhenDC3.

**Figure 5.**
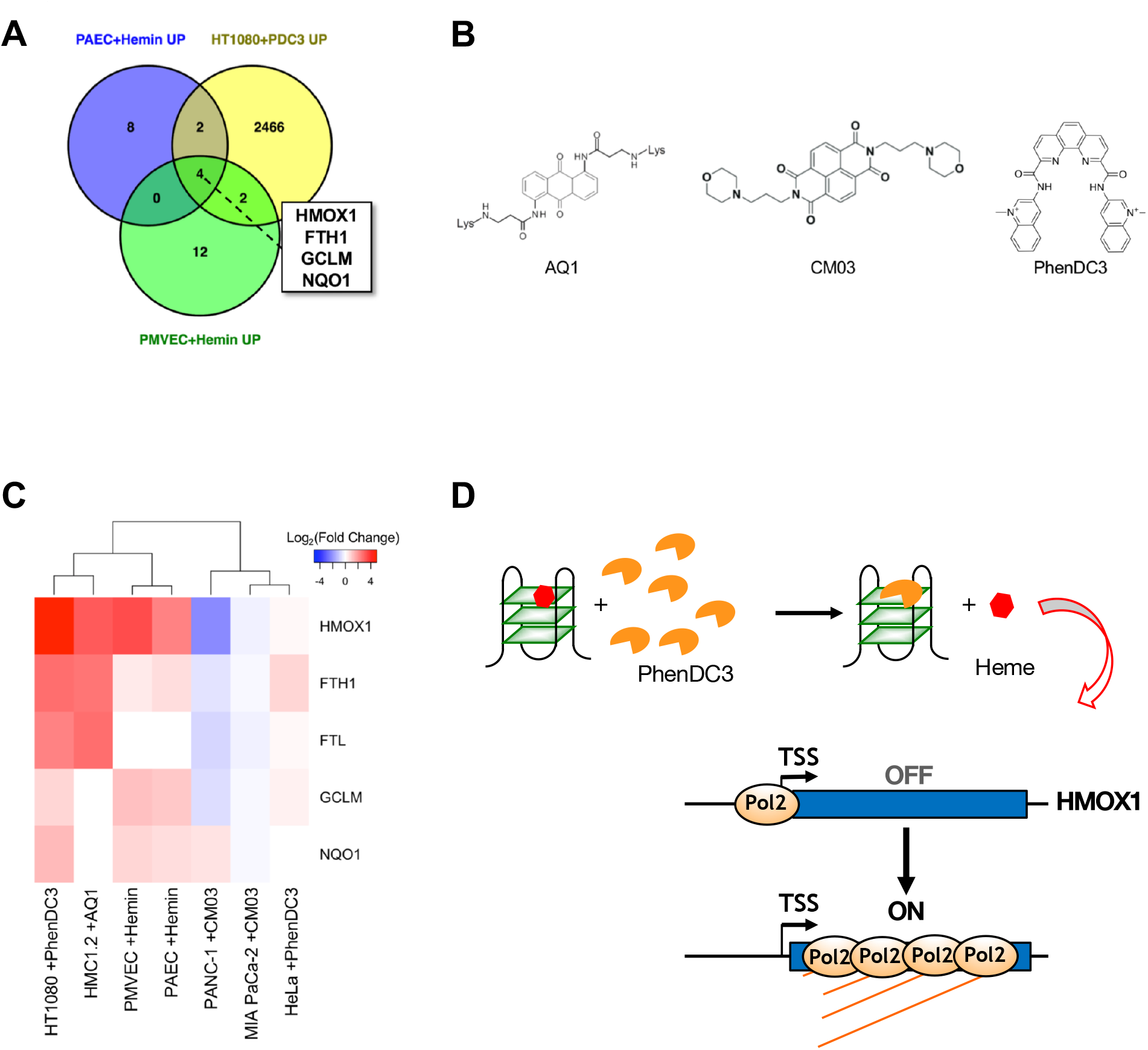
Common transcriptional targets of heme and synthetic G4 ligands. **(A)** Venn diagram of common genes significantly up-regulated (log_2_(Fold Change) > 0.5; FDR < 0.05) in PAEC cells treated with hemin, PMVEC cells treated with hemin and HT1080 cells treated with PhenDC3. **(B)** Structures of AQ1, CM03 and PhenDC3. **(C)** Hierarchical cluster analysis of log_2_(Fold Change) values for 5 heme metabolism related genes in datasets described in panel A. Red, log_2_FC >0; blue, log_2_FC <0. **(D)** Model for sequestration of heme by G-quadruplexes and induction of HMOX1 expression upon heme displacement. Regulation of HMOX1 is shown, but this model could be applicable to other NRF2 target genes.

To ask if synthetic G4 ligands other than PhenDC3 might induce transcription of heme-regulated genes, we took advantage of the three datasets reporting transcriptional response of human cells to synthetic G4 ligands that are currently publicly available [Halder R et al. 2012; Marchetti C et al. 2018; Zorzan E et al. 2018]. Two G4 ligands analyzed, AQ1 and CM03, are structurally distinct from one another and from PhenDC3 (**Figure 5B**). AQ1 is an anthraquinone derivative which can displace an end-stacking compound from quadruplexes but does not preferentially act at telomeric structures [Zorzan E et al. 2016], and which has been shown to affect proliferation of the mast cell leukeumia line HMC1.2 [Zorzan E et al. 2018]. CM03 is a trisubstituted naphthalene diimide derivative computer-designed to improve binding to telomeric quadruplexes and shown to function as a potent inhibitor of proliferation of pancreatic tumor cell lines and xenografts [Marchetti C et al. 2018]. Hierarchical cluster analysis showed clear overlap in transcriptional activation by PhenDC3, AQ1 and hemin, but not by CM03 (**Figure 5C**). This may in part reflect preferred binding of the synthetic ligands to distinct quadruplex structures.

These results support the hypothesis that heme is sequestered by G-quadruplexes, and can be displaced by G4 ligands to activate transcription of HMOX1 (**Figure 5D**) and other genes involved in heme and iron homeostasis. The ability of a G4 ligand to displace heme is likely to depend upon the structure of the ligand and its preferred quadruplex targets. Both DNA and RNA quadruplexes bind heme *in vitro*, and either could be the source of free heme released upon treatment of living cells with PhenDC3. Within a cell, interactions of heme with either DNA or RNA quadruplexes could result in local oxidation that compromises molecular integrity. In light of the robust catalytic activity that heme confers on quadruplex DNA and RNA aptamers, Sen and colleagues proposed that accumulation of transcripts bearing quadruplexes could be the source of potentially harmful 1- and 2-electron oxidative reactions [Poon LC et al 2011; Grigg JC et al. 2014]. Heme bound to RNA could contribute to the pathophysiology of repeat expansion diseases in which the transcript contains multimers of quadruplexes, such as the CGG repeat expansion in Fragile X; or sporadic cases of amyotrophic lateral sclerosis (ALS) and frontotemporal dementia (FTD) caused by an expansion of a GGGGCC repeat (reviewed in [Maizels N 2015]). Oxidation at heme bound to quadruplex DNA may also cause local damage, and could even constitute a potent local source of genomic instability. However, cells have evolved to repair isolated nicks and breaks in genomic DNA, and in an alternative view binding of DNA quadruplexes to heme might contribute to genomic stability, by capturing free heme near its site of origin and sequestering it until the quadruplex structure is resolved during replication or transcription. If so, sequestration of heme by quadruplexes represents an unanticipated role in maintenance of genomic stability, which may explain the abundance of G4 motifs and their dispersion throughout the genome.

Finally, the results reported here suggest that heme may function as an endogenous G4 ligand. If so, then changes in the levels of free heme in the nucleus may perturb expression not only of genes involved in heme and iron homeostasis, but also of other genes which are susceptible to perturbation by G4 ligands.

## Materials and methods

### Reagents for DNAzyme titration

4-(2-hydroxyethyl)-1-piperazineethanesulfonic acid (HEPES), 2,2’-azino-bis(3-ethylbenzothiazoline-6-sulfonic acid) (ABTS^2-^) diammonium salt, potassium acetate, DMSO, hemin and triton X-100 were purchased from Sigma-Aldrich. H2O2 30% was purchased from Merck. Inorganic salts and organic chemicals employed were high quality (analytical grade). Hemin and PhenDC3 were dissolved in DMSO to obtain two solutions at 1 mM concentration and stored in the dark at −20°C. ABTS^2-^ was freshly prepared in water to final concentration of 100 mM. Freshly prepared H2O2 was made on the spot by directly diluting the high concentration H2O2 30% (9.8 M) to the desired concentration (2 mM) for use. The reactions were carried out in 1 × K+ buffer (40 mM HEPES-NH_4_OH pH 7.8 with 20mM CH3COOK, 1% DMSO, and 0.05% Triton X-100), optimized to favor the highest activity of G4 DNAzymes (i.e. minimal background, with the complex-catalyzed reaction occurring 20 to 50 times faster than the background reaction), prepared by dilution of 10 × K+ buffer (400 mM HEPES-NH4OH, pH 7.8; 200 mM CH3COOK; 10% DMSO; 0.5% Triton x-100) made in ultrapure water and stored at 4°C before use. Assays used hemin 3 μM, ABTS^2-^ 500 μM, H_2_O_2_ 50μM, 3 μM G-quadruplexes (3 nmol); and PhenDC3 0, 0.5, 1, 3, 6 μM.

Sequences are shown in the table below for the PS2.M, PS5.M [Travascio P et al. 1998] and c-MYC [Simonsson T et al. 1998] DNA oligonucleotides used in this work. Oligonucleotides were purchased from Eurogentec (Liège, Belgique), dissolved in ultrapure water at room temperature and quantified by OD260nm using the molar extinction coefficient values provided by the manufacturer.

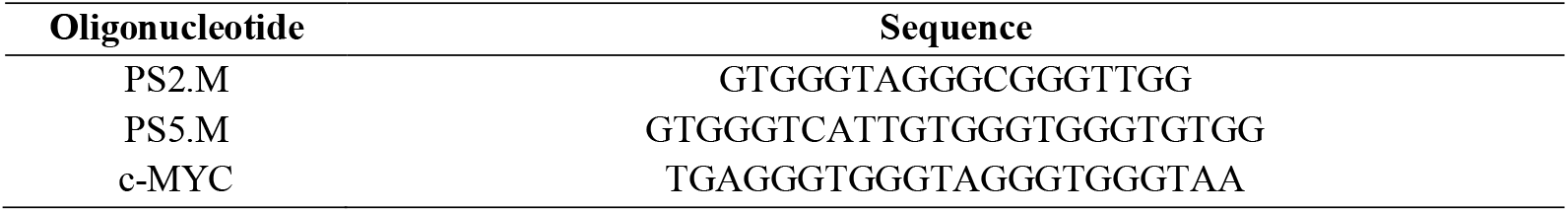

### Hemin/G-quadruplex DNA 1:1 complex titration in the presence of increasing concentration of PhenDC3

Absorption titrations using the chromogenic substrate ABTS^2-^ were performed on a Cary 300 UV-Vis spectrophotometer Agilent Technologies (Les Ulis, France), using a 1 cm path-length quartz cuvette. To prepare G4 DNA, the DNA oligos (300 μM in 1 × K^+^ buffer) were folded by heating at 95°C for 5 min and left to cool at room temperature over three hr. The DNA was distributed into separate tubes to prepare the sample for the titration experiments (final concentration 3 μM in 1 mL total volume for each sample). One-tenth volume 10 × K^+^ buffer (see above) was added to the DNA, for a final buffer concentration of 1 × K^+^ in ddH_2_O, and the samples were left to fold over 30 min. Hemin was added to a final concentration of 3 μM and the solutions were left to stand at room temperature for 30 min. Samples were brought to final concentrations of 0, 0.5, 1, 3, and 6 μM PhenDC3 by addition of appropriate volumes of 1 mM PhenDC3 in DMSO carrier. After 30 min incubation at room temperature, ABTS^2-^ was then added to each sample to a final concentration of 500 μM and the blanks were recorded with the spectrophotometer. To initiate the oxidation reaction, H_2_O_2_ was added to a final concentration of 50 μM, followed by quick mixing. Five min later the absorption spectra were recorded in the range of 400 to 800 nm. Final concentrations were as follows: 3 μM DNA, 3 μM Hemin, 1 × K^+^ (40 mM HEPES-NH_4_OH, pH 7.8; 20 mM CH_3_COOK; 1% DMSO; 0.05% Triton x-100), 0, 0.5, 1, 3, and 6 μM PhenDC3, 500 μM ABTS^2-^, and 50 μM H_2_O_2_ in a total volume of 1 mL.

### Docking of heme on the c-MYC DNA G-quadruplex

Receptor and ligand standard representation data files (Protein Data Bank, PDB format) were retrieved for the G-quadruplex structure of the c-MYC promoter (PDB accession: 2MGN [Chung WJ et al. 2014]) and the ligand Fe(III) protoporphyrin IX (PDB accession: HEM). We used a text editor to remove the HETATM residues from the receptor PDB file, as it contains information corresponding to the PhenDC3 ligand. We used the AutoDockTools (ADT) GUI v1.5.6 [Morris GM et al. 2009] to prepare the pdbqt files needed for docking, which contain atomic charges and atom type definitions.

We added hydrogen positions as well as topological information for the heme ligand to perform flexible docking by allowing 10 rotatable bonds (using the *TorsionTree* utility in ADT). Then, we defined a docking box covering the space to be searched around the receptor *(GridBox* utility in ADT), manually adjusting the size of the box and the exact position of its center before exporting it into a configuration file. We used AutoDock vina v1.1.2 [Trott O and Olson AJ 2010] for MacOSX to perform docking simulations from command line, increasing the exhaustiveness value from 8 (default) to 24, which gives a more consistent docking result [Forli S et al. 2016]. Nine docking poses were generated at the end of the run and ranked according to their binding energies, and these proved to be quite similar: all fell within the range of −6.9 to −5.2 kcal/mol (mean=-6.3±0.5 kcal/mol). Root-Mean-Square Deviation of atomic positions (RMSD) values were calculated relative to the best mode and are reported in the table below (l.b: lower-bound; u.b: upper-bound), showing one clear preferred pose relative to the receptor.

| Mode | Affinity<br>(kcal/mol) | Distance from best mode |  |
| --- | --- | --- | --- |
|  |  | RMSD l.b. | RMSD u.b. |
| 1 | -6.9 | 0 | 0 |
| 2 | -6.8 | 0.655 | 5.719 |
| 3 | -6.8 | 1.318 | 5.862 |
| 4 | -6.5 | 1.272 | 1.663 |
| 5 | -6.3 | 4.079 | 7.779 |
| 6 | -6.0 | 2.242 | 3.312 |
| 7 | -5.9 | 2.403 | 6.832 |
| 8 | -5.7 | 4.017 | 9.427 |
| 9 | -5.6 | 4.704 | 9.914 |

Visualization of the docked receptor-ligand structures was performed using PyMOL v2.3.1 for MacOSX [The PyMOL Molecular Graphics System, Version 2.0 Schrödinger, LLC]. All ligands poses were located in the same region of the G-quadruplex, from visual inspection. Finally, visualization of the G4 structure bound to PhenDC3 was also performed on PyMOL, using the original 2MGN PDB file.

### Cell culture and cell viability, cell death and cell cycle assays

HT1080 fibrosarcoma cells were cultured in Dulbecco’s Modified Essential Medium (DMEM) with 5% fetal bovine serum, glutamate, penicillin, and streptomycin. To assess cell viability, a 96 well plate was seeded with 2000 cells per well and cultured for 24 hr at 37°C in 5% CO2. Samples were then brought to final concentrations of 1% DMSO and 0, 10 or 20 μM PhenDC3 by addition of DMSO carrier alone or 1 mM PhenDC3 in DMSO. After an additional 24 or 48 hr culture, cell viability was assayed by addition of CellTiter Glo (Promega) flowed by quantification of luminescence using a Fluostar Omega (BMG Labtech). Cell death was measured by staining cells with LIVE/DEAD Fixable Near-IR Dead Cell Stain (Invitrogen) for 30 min, resuspending them in 3.7% formaldehyde, and analysis using a FACS Canto II with 633 nm excitation and 780 nm emission. Cell cycle profiles of DAPI-stained cells were generated by flow cytometry on a LSRII with 355 nm excitation and 450 nm emission. Results of LIVE/DEAD assays and cell cycle profiles were analyzed using FlowJo v10.

We chose to examine cells treated with 20 μM PhenDC3 for 48 hr based on control experiments which showed that effects of treatment were barely perceptible at 24 hr, and that 20 μM PhenDC3 was relatively non-toxic (>90% viable; **Figures 3; S1**). Cells were split 24 hr prior to treatment and cultured in six plates at 3×105 cells per 10 cm plate, in 10 ml of media per plate. Treatment was initiated by replacing culture media with 10 ml of fresh media containing either 100 μl of 2 mM PhenDC3 stock in DMSO (20 μM final concentration; 3 culture plates), or DMSO only (3 culture plates). At 48 hr, cells were removed from each plate by trypsinization and the resulting six samples separately processed for RNA-seq.

### RNA extraction and sequencing

Cells were pelleted by centrifugation, re-suspended in TRIzol Reagent (Life Technologies), and total RNA was harvested using the supplier’s protocol and stored at −80°C. RNA concentration was adjusted to 0.5 μg/μl, and 5 μg of each sample was submitted to the UW High-Throughput Genomics Center (htSeq) for library preparation and sequencing. RNA samples were tested for quality using a Bioanalyzer (Agilent Technologies). mRNA library preparation was carried out using poly-dT selection. After library construction, multiplexed libraries were sequenced on an Illumina Hi-Seq to 36 bp in paired-end mode. Data were deposited in the Gene Expression Omnibus (GEO) database, accession number GSE60630.

### RNA-seq data analysis

Reads were aligned to the human reference genome hg19 using STAR v2.7 [Dobin A et al. 2013] with the ENCODE processing pipeline standard settings for RNA-seq data (--*outFilterType* BySJout, --*outFilterMultimapNmax* 20, --*alignSJoverhangMin* 8, --*alignSJDBoverhangMin* 1, --*outFilterMismatchNmax* 999, --*alignIntronMin* 20, --*alignIntronMax* 1000000 and --*alignMatesGapMax* 1000000). Reads mapping uniquely to GENCODE-annotated genes were summarised using *featureCounts* [Liao Y et al. 2014]. The raw gene count matrix was imported into the R environment [R Core Team 2018] for further processing and analysis. Genes with low read counts (less than ~10 reads in more than 3 samples) were filtered out, leaving a set of 20,000 genes to test for differential expression between control (DMSO) and treated (+PhenDC3) conditions. Differential expression analysis was carried out using the R package *DESeq2* version 1.18.1 [Love MI et al. 2014]. Unless otherwise stated, differentially expressed genes were identified based on false discovery rate (FDR, using Benjamini-Hochberg adjusted *p*-values), with the threshold FDR < 1%. A fold-change cutoff was used to create the sets of genes used for G4 enrichment analysis: only ≥2-fold (≥1-log_2_fold) differentially expressed genes (2xDEGs) were retained. Hierarchical clustering was performed for the top 500 differentially expressed genes using Euclidean distances. The *cuffdiff* tool from Cufflinks v2.1.1 [Trapnell C et al. 2013] was used to calculate differentially expressed isoforms with the default q-value cutoff of < 0.05. Statistics for isoform expression levels and differential isoform expression are available in the GEO repository (GSE60630).

### G4 motif enrichment analysis

G4 motifs were defined as carrying 4 or more runs of 3 or more Gs, separated by 1-12 nt loops, and were located using regular expression matching as previously described [Gray LT et al. 2014]. Results were stored as bed files with coordinates as well as wig files for visualization and further processing. G4 enrichment was assessed using R scripts that calculated the frequency of G4 overlap at each position and on each strand near the TSS (+/- 250 bp) and near the 5’ end of first intron (1-250 bp). TSS and intron locations were retrieved from the Ensembl genes table in the UCSC hg19 iGenomes database using gene symbols. Briefly, we determined false discovery rates (FDR) by selecting 1000 sets of genes of matching size (e.g. for the 716 genes up-regulated >2-fold, we assessed 1000 sets of 716 random genes) from the pool of all reliably detected genes, which were subjected to the same count analysis described above. G4 motif enrichment or depletion found in fewer than 5% of the randomly selected sets was considered significant (FDR < 0.05). R scripts were used to tabulate and assemble results.

### Gene Set Enrichment Analysis (GSEA) and protein-protein interaction network

The REACTOME pathways enrichment analysis was done using the GSEA (Gene Set Enrichment Analysis) software [Subramanian A et al. 2005] with the Molecular Signatures Database collection on the identified up-regulated gene lists. Significantly enriched pathways were selected based on FDR q-values (P-value adjustment for multiple hypergeometric tests), with the cutoff FDR<5%. The NetworkAnalyst platform [Xia J et al. 2015] and the Cytoscape 3.5 software [Shannon P et al. 2003] were used to create and visualize the first-order (proteins that directly interact with a given protein) protein-protein interactions based on the high-confidence STRING interactome information [Szklarczyk D et al. 2015]. High-confidence interactions were defined has having a confidence score > 0.9 and experimental evidence. As the first-order network was very dense, we constructed a minimally connected network containing all the seed genes as well as essential non-seed interactors that maintain the network connection. To do this, we compute pair-wise shortest paths between all seed nodes, and remove the nodes that are not on the shortest paths. The obtained minimum first-order network consisted of 1,301 nodes and 4,038 edges, with larger nodes indicating higher connectivity.

## Acknowledgments

### Funding

This work was supported by the US National Institutes of Health National Cancer Institute (P01CA077852 to N.M.), by the Agence Nationale de la Recherche (ANR 14-CE35-0003-02 to A.N. and M.P.-T.F.) and by the PIC3i program from Institut Curie (n°91730 “Prospects of Anticancer” to A.N., M.P.-T.F. and A.L.V.). E.P.L. is a recipient of a doctoral fellowship from the French Ministry of Education, Research and Technology.

### Authors’ contributions

L.T.G. carried out cell culture, PhenDC3 treatment and prepared RNA. L.T.G. and E.P.L. designed the *in silico* analytical framework and wrote the code for analyzing and presenting the data. E.P.L. performed the final version of the RNA-seq data analysis and implemented the docking simulations. D.V performed the hemin/G-quadruplex DNAzyme assay. C.G. synthesized PhenDC3. A.N., M.P.-T.F., A.L.V. and N.M. conceived of and supervised the project. The manuscript was written by L.T.G., E.P.L. and N.M. All authors read and approved the final draft of the manuscript.

