## Supplementary Figures for "G-quadruplexes sequester free heme in living cells"

Departments of [1]Biochemistry and [5]Immunology, University of Washington, 1959 NE Pacific St., Seattle, WA 98195, USA

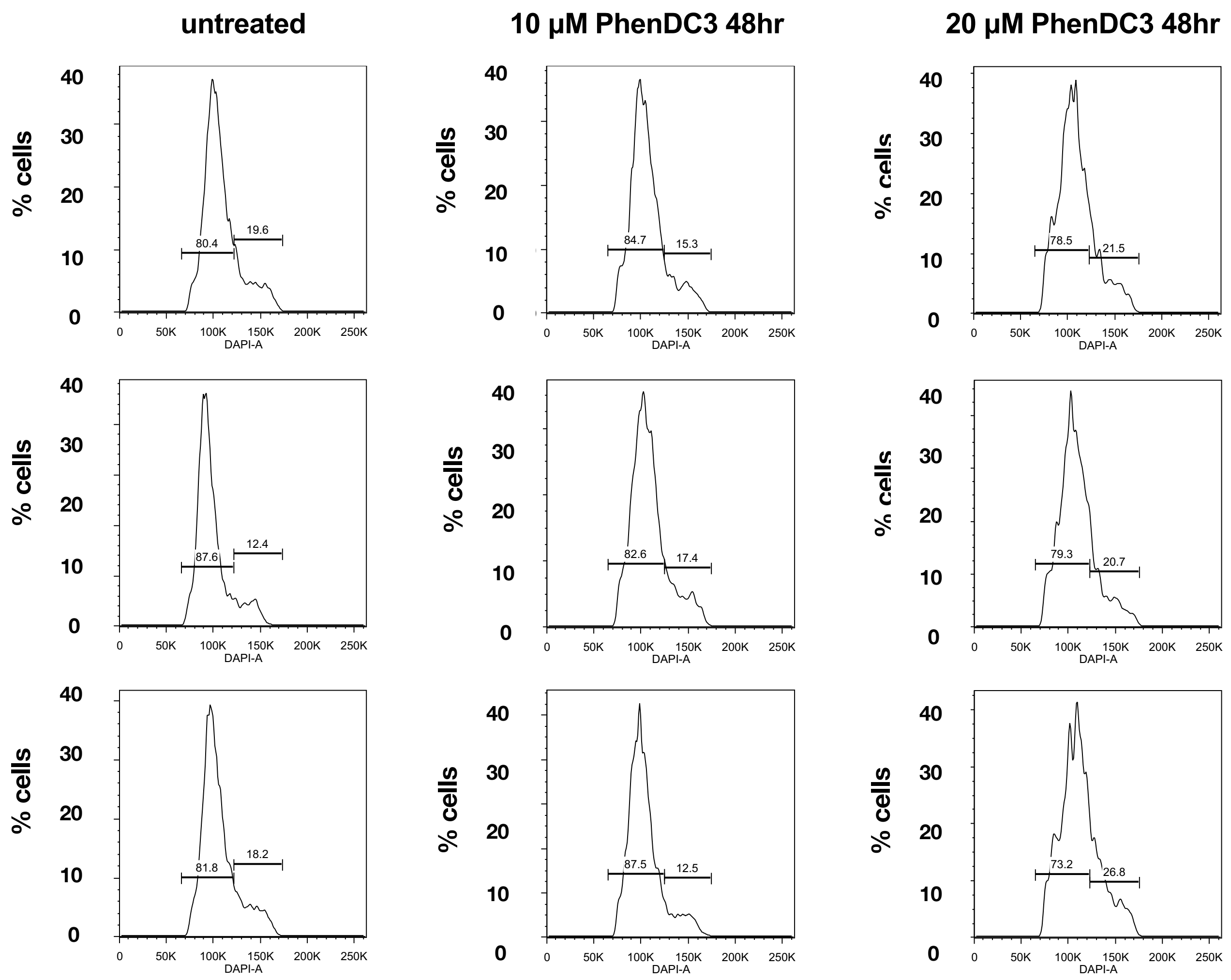


**Figure S1. Effects of PhenDC3 on cell viability and cell cycle.**

DAPI cell cycle profiles for populations treated with 0, 10, or 20 µM PhenDC3 for 48 hr. The triplicate replicates analyzed by RNA-Seq included the two samples treated with 20 µM PhenDC3 shown here, and a third sample shown in Figure 2B.

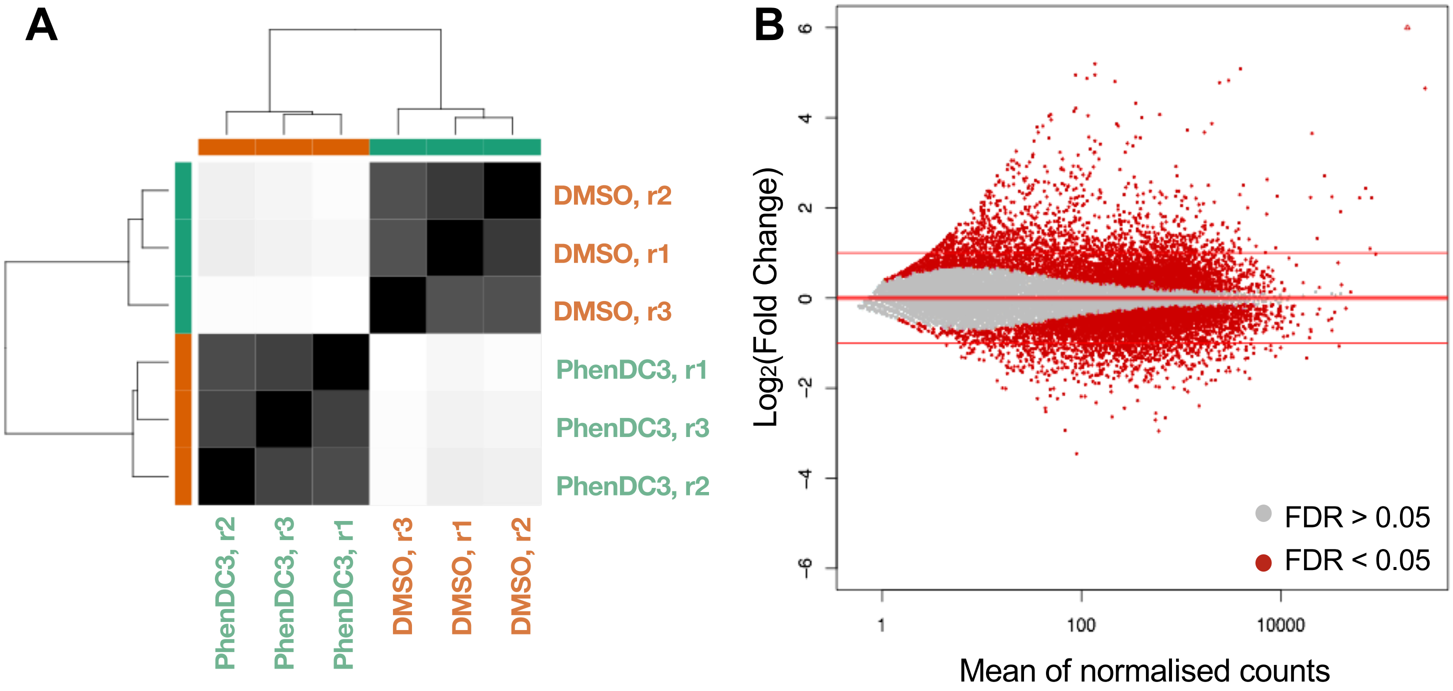


**Figure S2. PhenDC3 treatment induces significant transcriptional changes in HT1080 fibrosarcoma cells.**

(A) Euclidean sample-to-sample distances calculated from the normalized count data of HT1080 cells treated with 20 µM PhenDC3 for 48 hours. Hierarchical clustering of distance matrices shows a clear distinction between untreated (DMSO) and treated (+PhenDC3) samples. (B) MA plot showing differentially expressed genes between untreated and treated samples. Each dot represents one gene; red, genes with significantly altered expresssion values (FDR < 0.05); gray, genes without significantly altered expresssion values (FDR > 0.05).

| Gene Set | Strand | TSS -250 nt | TSS +250 nt | 5' end of intron 1 +250 nt |
|---|---|---|---|---|
| Up-regulated (N = 1,242) | (+) | Enriched (<0.001) | Enriched (<0.001) | Enriched (<0.001) |
| | (-) | Enriched (<0.001) | Enriched (<0.001) | Enriched (<0.001) |
| Down-regulated (N = 970) | (+) | n.s. | n.s. | Depleted (0.002) |
| | (-) | Depleted (0.013) | n.s. | n.s. |

**Figure S3. G4 motif frequency for differentially expressed genes in HeLa cells treated with PhenDC3.**

G4 motif enrichment and depletion statistics for genes up and down-regulated by PhenDC3 treatment of HeLa cells. Significantly enriched (green); depleted (red); FDR > 5% (0.05) is considered non-significant (n.s., no highlight). Gene lists were retrieved from supplementary data from Halder R et al. *BMC Res Notes* 5, 138 (2012).

A

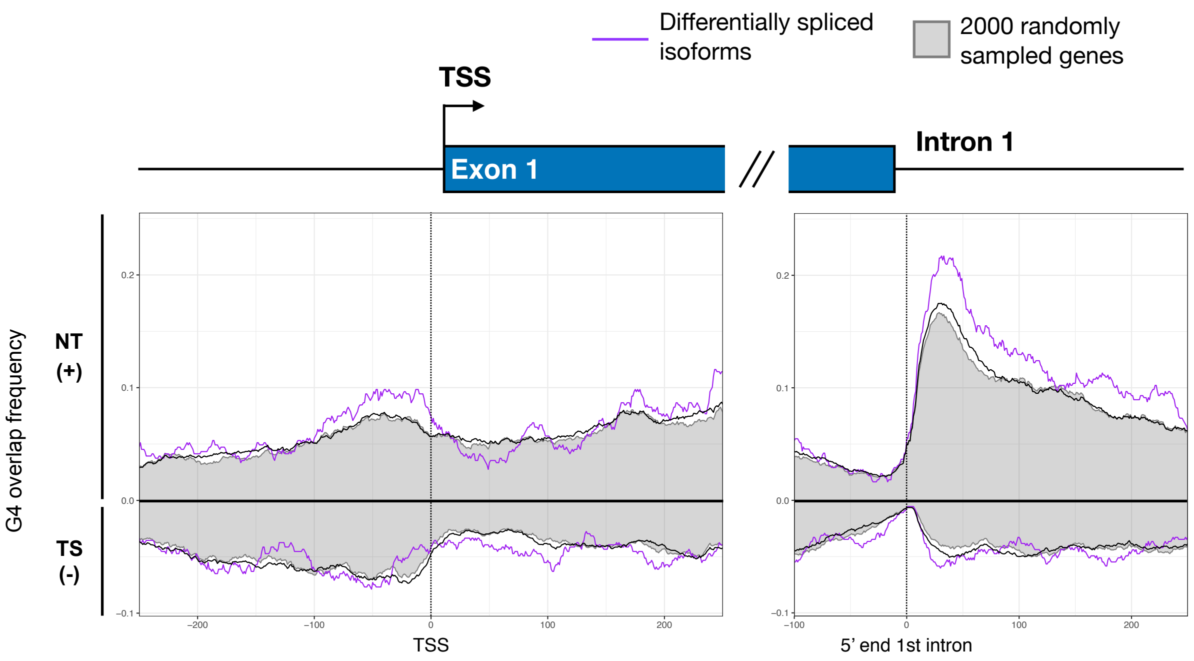


B

| Isoform Set | Strand | TSS | | 5' end of intron 1 |
|---|---|---|---|---|
| | | -250 nt | +250 nt | +250 nt |
| Differentially spliced isoforms | (+) | Enriched (0.025) | n.s | Enriched (0.004) |
| | (-) | n.s | n.s | n.s |

**Figure S4. Quadruplexes at the 5’ end of intron 1 correlate with altered splicing in PhenDC3-treated cells. (A)** G4 motif overlap frequency near the TSS and downstream of the 5’ end of the first intron of isoforms that exhibited altered splicing in response to PhenDC3 (n = 415; lavender line or splicing not significant; black line); and for 2000 randomly selected isoforms (gray). Frequency is calculated as the percent of genes that have a G4 motif on the specified strand at each position. **(B)** G4 enrichment and depletion statistics for 250 nt windows flanking the TSS and the 5’ end of the first intron, relative to 1000 sets of randomly selected genes drawn from a pool of all genes expressed at detectable levels. FDR values in parentheses. Significantly enriched (green); FDR > 5% (0.05) is considered non-significant (n.s., no highlight).

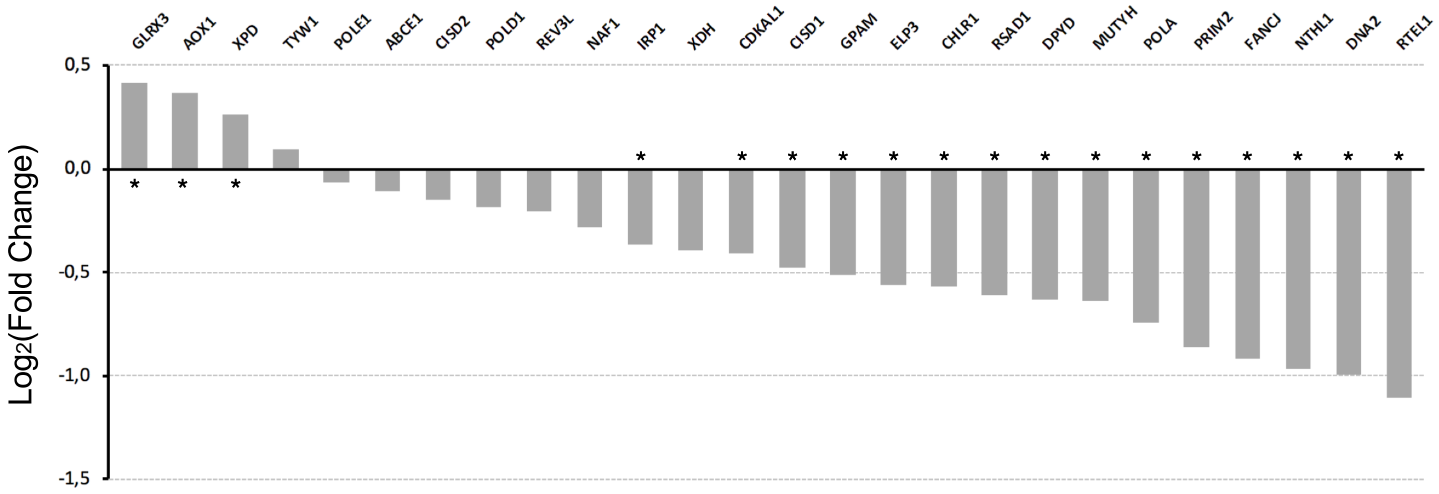


**Figure S5. Response of iron-sulfur proteins to PhenDC3 treatment.**

$Log_2$ fold changes for genes encoding iron-sulfur proteins. *, FDR < 0.05.
